# m6A-TSHub: unveiling the context-specific m6A methylation and m6A-affecting mutations in 23 human tissues

**DOI:** 10.1101/2022.01.12.476117

**Authors:** Bowen Song, Daiyun Huang, Yuxin Zhang, Zhen Wei, Jionglong Su, João Pedro de Magalhães, Daniel J. Rigden, Jia Meng, Kunqi Chen

**Author notes:** To whom correspondence should be addressed (Chen K) and (Huang D).

## Abstract

As the most pervasive epigenetic marker present on mRNA and lncRNA, *N^6^*-methyladenosine (m^6^A) RNA methylation has been shown to participate in essential biological processes. Recent studies revealed the distinct patterns of m^6^A methylome across human tissues, and a major challenge remains in elucidating the tissue-specific presence and circuitry of m^6^A methylation. We present here a comprehensive online platform m6A-TSHub for unveiling the context-specific m^6^A methylation and genetic mutations that potentially regulate m^6^A epigenetic mark. m6A-TSHub consists of four core components, including (1) m6A-TSDB: a comprehensive database of 184,554 functionally annotated m^6^A sites derived from 23 human tissues and 499,369 m^6^A sites from 25 tumor conditions, respectively; (2) m6A-TSFinder: a web server for high-accuracy prediction of m^6^A methylation sites within a specific tissue from RNA sequences, which was constructed using multi-instance deep neural networks with gated attention; (3) m6A-TSVar: a web server for assessing the impact of genetic variants on tissue-specific m^6^A RNA modification; and (4) m6A-CAVar: a database of 587,983 TCGA cancer mutations (derived from 27 cancer types) that were predicted to affect m^6^A modifications in the primary tissue of cancers. The database should make a useful resource for studying the m^6^A methylome and genetic factor of epitranscriptome disturbance in a specific tissue (or cancer type). m6A-TSHub is accessible at: www.xjtlu.edu.cn/biologicalsciences/m6ats.

## Introduction

Among the more than 150 distinct chemical modifications naturally decorating cellular RNAs [1], *N^6^*-methyladenosine (m^6^A) is the most pervasive marker present on mRNA and lncRNA, and has been associated with a number of essential biological functions and processes [2, 3], including mRNA stability [4], splicing [5], translation [6, 7], heat shock [8], DNA damage [9], and embryonic development [10]. Increasing evidence has indicated a critical role of m^6^A dysregulation in various human diseases, especially multiple cancers, such as breast cancer [11, 12] and prostate cancer [13]. For example, inhibition of an m^6^A methyltransferase (METTL13) could be used as a potential therapeutic strategy against acute myeloid leukemia [14].

Developed in 2012, m^6^A-seq (MeRIP-seq) was the first whole transcriptome m^6^A profiling approach [15, 16]. It relies on antibody-based enrichment of the m^6^A signal, enabling identification of m^6^A-containing regions with a resolution of around 100nt. Currently, m^6^A-seq is still the most popular m^6^A profiling approach and has been applied in more than 30 different organisms. Besides m^6^A-seq, recent advances in integration of UV cross-linking, enzymatic activity and domain fusion have offered improved even base-resolution m^6^A detection through techniques such as, miCLIP/m^6^A-CLIP-seq [17, 18], m^6^A-REF-seq [19] and DART-seq [20]. However, compared with m^6^A-seq, these approaches require more complicated experimental procedures, and have therefore been applied in fewer biological contexts.

To date, more than 120 computational approaches have been developed for the computational identification of RNA modifications [21, 22] from the primary RNA sequences. These include the iRNA toolkits [23–31], MultiRM [32], DeepPromise [22], RNAm5CPred [33], SRAMP [11], Gene2vec [34], PEA [35], PPUS [36], WHISTLE [37], m5UPred [38], WeakRM frameworks [39, 40], m6ABoost [41], PULSE [42], m6AmPred [43], BERMP [44] and MASS [45]. Together, these efforts greatly advanced our understanding of multiple RNA modifications at different RNA regions and in various species (see recent reviews [22, 46–48]). A number of epitranscriptome databases have been constructed. MODOMICS collects the pathways related to more than 150 different RNA modifications [1]. RMBase [49], MeT-DB [50] and m^6^A-Atlas [51] assembled millions of experimentally validated m^6^A sites. REPIC was established as a comprehensive atlas for exploring the association between m^6^A RNA methylation and chromatin modifications [52]. ConsRM provides the conservation score of individual m^6^A sites at base-resolution, which can be used to differentiate the functionally important and ‘passenger’ m^6^A sites [53]. M6A2Target compiled the target molecules of m^6^A methyltransferases, demethylases and binding proteins [54]. This work has extended our knowledge of the functional epitranscriptome, and greatly facilitated relevant research. Special efforts have also been made to explore the effects of genetic variants on RNA modifications and their association with various diseases. m^6^AVar [55] was the first database that focused on the genetic factors related to epitranscriptome disturbance. It documented more than 400,000 m^6^A-affecting genetic variants, which were further labeled with disease and phenotype associations identified from GWAS analysis. This prediction framework was improved and later applied to eight other RNA modifications (m^5^C, m^1^A, m^5^U, Ψ, m^6^Am, m^7^G, and 2’-O-Me, and A-to-I) by RMVar [56] and RMDisease [57]. These above databases systematically revealed the general association between epitranscriptome layer dysregulation and various diseases (see a recent review [58]).

Existing computational approaches for epitranscriptome analysis have been quite successful in providing lots of useful information; however, most of them failed to consider the tissue-specificity of m^6^A epitranscriptome [59, 60]. Indeed, recent study by *Liu et al*., unveiled distinct tissue-specific signatures of the m^6^A epitranscriptome in human and mouse [61], which are induced by context-specific expression of m^6^A regulators (methyltransferases, demethylases and RNA binding proteins) [62] and genetic drivers [63]. Nevertheless, most existing approaches for RNA modification sites prediction completely ignore the context-specificity of the epitranscriptome and simply assume a single model for different tissues, undermining their accuracy and applicability. To the best of our knowledge, the only three approaches that clearly support the identification of tissue-specific m^6^A methylation are im6A-TS-CNN [64], iRNA-m6A [65], and TS-m6A-DL [66], all covering only three human tissue types (brain, liver and heart). Similarly, when screening for the genetic variants that can affect RNA modifications, previous work assumes a consistent influence in different tissues (see Table S1 for a detailed description and comparison). However, since different epitranscriptome patterns were observed among different tissues, genetic mutations that can alter m^6^A methylation in one tissue may not necessarily function similarly in a different tissue. Likewise, there are significant differences in incidence, mortality and molecular signatures across cancer originating from different tissues [67, 68]. It is therefore highly desirable to develop approaches that could take full advantage of the tissue-specific RNA methylation profiles so as to make more reliable predictions with respect to a specific tissue type [69]. And this is particularly critical for studying the epitranscriptome circuits of diseases that are explicitly associated with a specific tissue, such as, cancers.

To address this issue, we present here a comprehensive online platform m6A-TSHub for unveiling the context-specific m^6^A methylation and m^6^A-affecting mutations in 23 human tissues. m6A-TSHub consists of four core components:

i. m6A-TSDB: a database for 184,554 experimentally validated m^6^A-containing peaks (m^6^A sites) derived from 23 distinct human normal tissues and 499,369 m^6^A-containing peaks (m^6^A sites) from 25 matched tumor conditions, extracted from 233 m^6^A-seq samples, respectively.
ii. m6A-TSFinder: an integrated online server for the prediction of tissue-specific m^6^A modifications in 23 human tissues, built upon a gated attention based multi-instance deep neural network.
iii. m6A-TSVar: a web server for systemically assessing the tissue-specific impact of genetic variants on m^6^A RNA modification in 23 human tissues.
iv. m6A-CAVar: a database of 587,983 TCGA cancer mutations (derived from 27 cancer types) that may lead to the gain or loss of m^6^A sites in the corresponding cancer-originating tissues.

In addition, the m^6^A-associated variants were also annotated with their potential post-transcriptional regulatory roles, including RBP binding regions, microRNA targets, and splicing sites, along with their known disease and phenotype linkage integrated from GWAS Catalog [70] and ClinVar databases [71]. The m6A-TSHub is freely accessible at: www.xjtlu.edu.cn/biologicalsciences/m6ats, and should be a useful resource for studying the m^6^A methylome and genetic basis of epitranscriptome disturbance with respect to a specific cancer type or tissue. The overall design of m6A-TSHub is shown in Figure 1.

**Figure 1.**
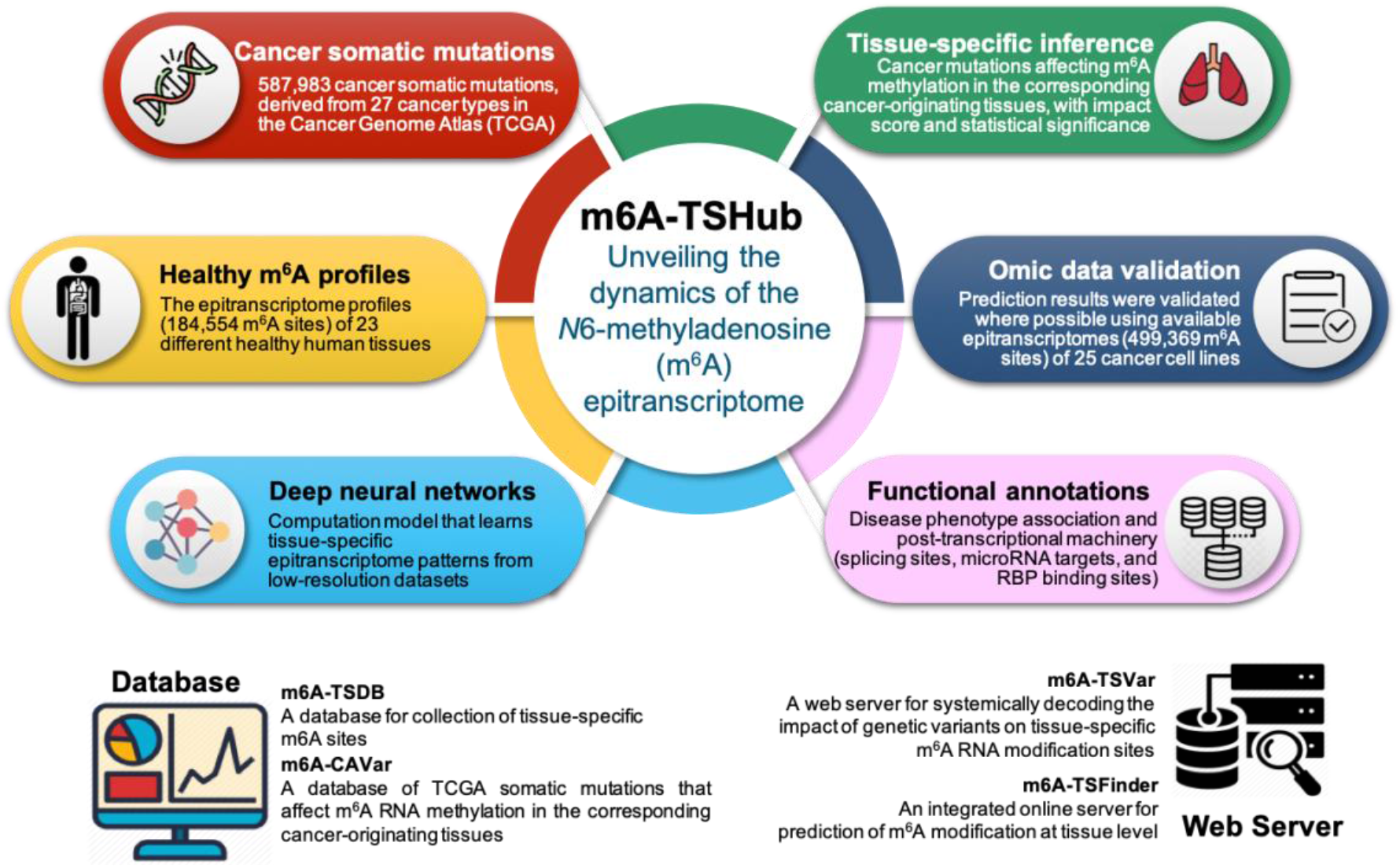
The overall design of m6A-TSHub. By integrating 184,554 m^6^A sites detected from 23 different healthy human tissues (m6A-TSDB), a deep learning framework that learns tissue-specific RNA methylation patterns was developed (m6A-TSFinder). The effect of genetic variants on tissue-specific m6A sites was then evaluated (m6A-TSVar). A total of 587,983 cancer somatic mutations were predicted to be able to affect m^6^A methylation of RNA in their corresponding cancer-originating tissues. The predicted m^6^A-affecting SNPs were then systematically validated using available cancer epitranscriptome datasets, and then functionally annotated with disease and phenotype association from genome-wide association studies (GWAS), along features relating to the post-transcriptional machinery (microRNA target sites, splicing sites and RNA binding protein binding sites) that are potentially mediated by m^6^A modification (m6A-CAVar). A web interface was constructed to enable the exploration, query, online analysis, and download of relevant results and data.

## Data collection and processing

### Data resource (m6A-TSDB)

We collected the epitranscriptome profiles of 23 healthy human tissues, from which the tissue-specific RNA methylation patterns were learned using deep neural networks. Specifically, the raw sequencing data of 78 m^6^A-seq samples were downloaded directly from NCBI GEO [72] and National Genomics Data Center [73] (Sheet S1). Adaptors and low quality nucleotides were removed by Trim Galore [74], followed by quality control using FastQC. The processed reads were then aligned to the reference genome hg19 by HISAT2 [75]. The m^6^A enriched regions (peaks) located on transcripts were detected by exomePeak2 [76] using its default setting with GC contents corrected. In total, m^6^A profiling samples from 23 human healthy tissues (184,554 m^6^A-containing peaks) were processed, we filtered all obtained m6A enriched regions to remain peaks with at least one DRACH consensus motif located, and using these peak regions containing tissue-specific m^6^A signal as positive data. Negative data was randomly collected from non-peak regions located on the same transcript of the corresponding positive data, and cropped to balance the length and number between positive and negative regions (with a positive to negative ratio of 1:1). The genomic sequences of both positive and negative regions were then extracted for developing the tissue-specific m^6^A prediction model.

To evaluate the effect of cancer somatic variants on m^6^A methylation in their originating tissues, a total of 2,587,191 cancer somatic variants from 27 different cancer types were obtained from The Cancer Genome Atlas (TCGA) (release version v27.0-fix) [77] (Sheet S2). Meanwhile, 155 m^6^A-seq samples profiling the epitranscriptome (499,369 m^6^A-containing peaks) of 25 cancer cell lines (corresponding to 17 tissue types) were also obtained using the same data processing pipeline (Sheet S1), which were used for the validation of the predicted effects on m^6^A methylation of the variants (detailed in the following).

### Learning tissue-specific m^6^A methylation with deep neural networks (m6A-TSFinder)

The purpose of weakly supervised learning is to develop predictive models by learning from weakly labeled data, such as m^6^A peaks of low resolution detected by the m6A-seq (or MeRIP-seq) technique [15, 16]. Unlike supervised learning based on single-nucleotide resolution data, it works for the case where only coarse-grained labels (indicating whether a genome bin contains a m^6^A site) are available for these peaks of various lengths. We previously proposed a general weakly supervised learning framework WeakRM [78], which takes labels at the sequence level (rather than a nucleotide level) as input and predicts the sub-regions that are most likely to contain the RNA modification. As a simplified illustration showed in Figure 2, the m6A-TSFinder framework is divided into several sub-sections. Firstly, multi-instance learning treats each entire RNA sequence as a ‘bag’, with multiple ‘instances’ within the ‘bag’ determined by a fixed-length sliding window. Previous studies have shown that a 40-50nt context region is sufficient for modification predictions. Therefore, in m6A-TSFinder, a sliding window of 50-nt was used, which is also helpful to improve the prediction resolution. Secondly, the RNA instances were fed into m6A-TSFinder model using one-hot encoding, which is widely used in deep learning-based models. The extracted instances pass through the same feature extraction module (the weights of the network are shared in this module) and output instance-level features. The network architecture of the feature extraction section used in m6A-TSFinder includes: the first convolutional layer to capture motifs; a max-pooling layer to remove weak features and enlarge the receptive field; a dropout layer that prevents overfitting in training, and a second convolutional layer which learns local dependencies among motifs. In order to further improve the performance of the model, in m6A-TSFinder, we use a long short-term memory (LSTM) layer to replace the second convolutional layer, so that the model can learn the long-range dependence of the motif while maintaining local dependence. Lastly, gated attention was used as the score function to obtain bag-level probabilities from multiple instance-level features. The gated attention module consists of three fully connected layers. The first two layers learn hidden representations from the instance features using tanh and sigmoid activation functions. Their element-wise multiplication is then sent to the third fully connected layer, which learns the similarity between the product and a context feature vector and outputs an attention score for each instance. The score is further normalized using the softmax function, so that the weights of all instances add up to 1. The weighted summation of instance features is treated as the bag-level feature and used to output the final probability score. Together, our model can be trained end-to-end using the binary cross-entropy loss calculated by the bag-level label. Our model was trained using the Adam optimizer under the Tensorflow framework. The learning rate was initially set to 1e-4, and gradually decayed to 1e-5 during the training process of 20 epochs. It is worth mentioning, when the number of instances is consistently set to 1, the weight of the instance is always 1, and the label becomes the instance level. In that case, the gated attention module is degraded, and the network becomes a strong supervised learning framework with two feature extraction layers.

**Figure 2.**
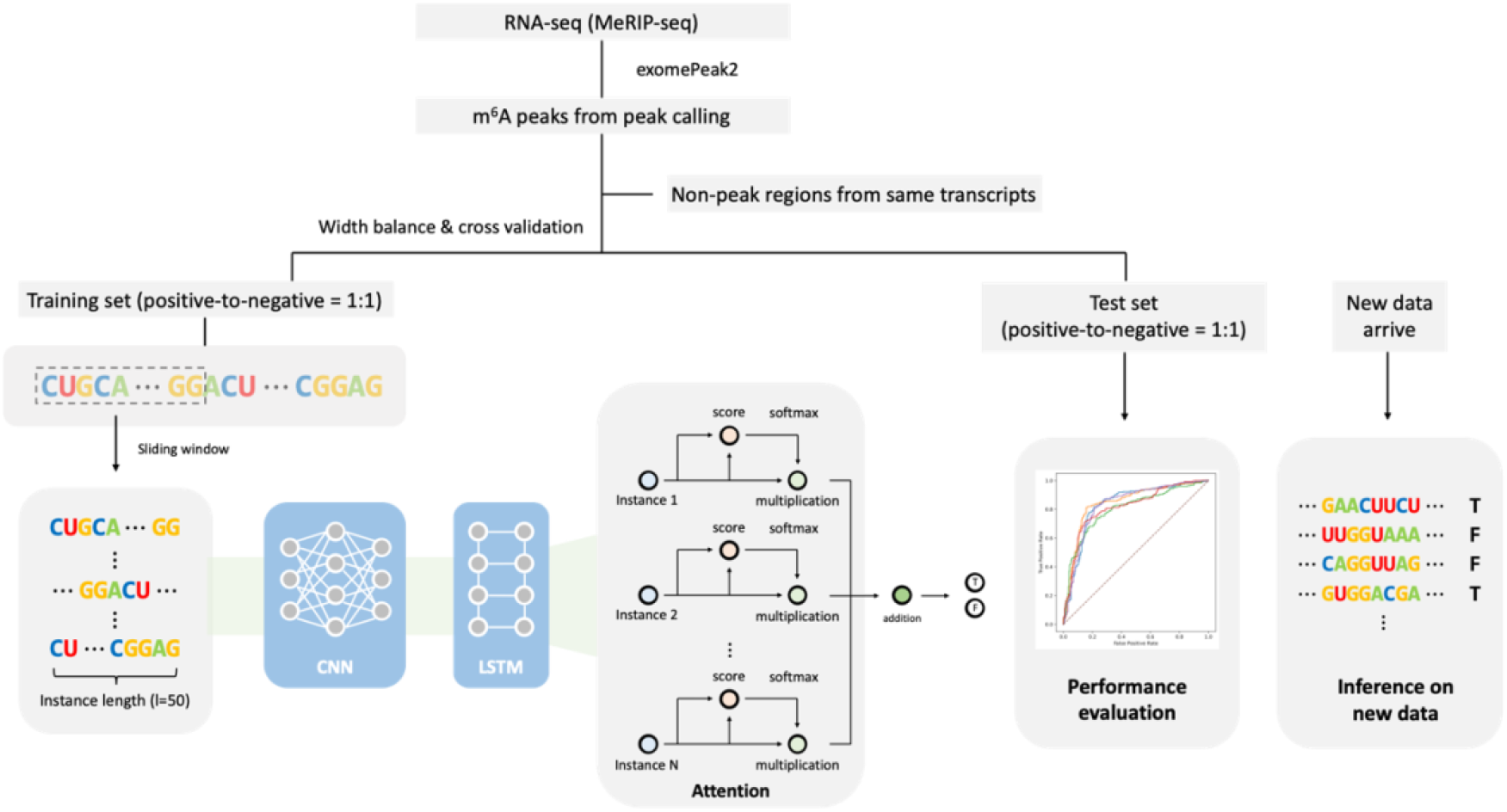
A simplified graphic illustration of the proposed m6A-TSFinder framework.

### Decoding the tissue-specific effect of variants on m^6^A methylation (m6A-TSVar & m6A-CAVar)

Similar to previous studies [55, 56, 79, 80], a cancer somatic variant is defined as tissue-specific m^6^A variant if it could lead to the gain or loss of m^6^A methylation in a specific tissue. The tissue-specific inference was made possible by our deep neural network model m6A-TSFinder. Specifically, the predicted tissue-specific m^6^A variants were further classified into 3 confidence levels - low, medium and high (see Figure 3).

**Figure 3.**
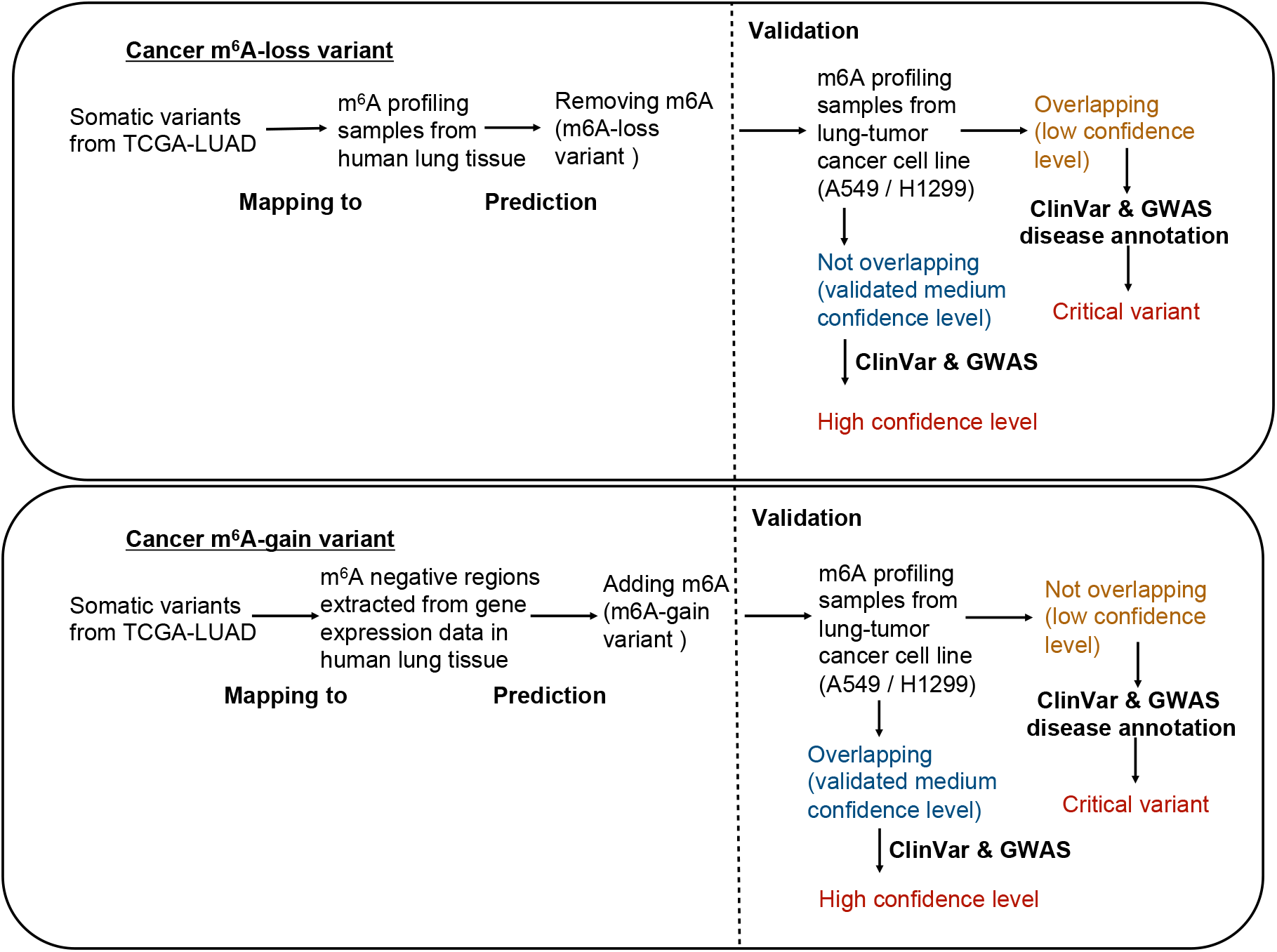
Workflow of how to determine the confidence level of m^6^A variants. Three types of confidence levels were applied. The cancer-driving somatic variants were extracted from TCGA-projects, and mapped to the m^6^A profiling samples derived from corresponding tumor-growth tissue. A tissue-specific weakly supervised model was then applied to obtain m^6^A-associated variants labeled in low confidence level. m^6^A profiling samples from tumor-growth tissues were then used for validation of the prediction results, the validated portion was classified into medium confidence level. Lastly, all variants with medium confidence level were annotated with disease information from ClinVar and GWAS, and then classified into the high confidence group. Lung tissue, healthy and cancerous, is used as an example here, the same protocol was followed for all 23 tissues.

#### Low confidence level

An m^6^A-associated variant with a low confidence level was defined directly by the tissue-specific prediction model. For example, a synonymous somatic variant (chr5:92929473, positive strand, C>T, TCGA barcode: TCGA-49-6742-01A-11D-1855-08) was extracted from TCGA-LUAD project, which was then predicted to eliminate the methylation of an experimentally validated m^6^A-containing region (chr5:92929314-92929786, positive strand) originally detected in human lung tissue [61].

#### Medium confidence level

m^6^A variants of medium confidence level are those that can be verified on available epitranscriptome data from cancer samples originated from the matched tissue. Follow the low confidence level mentioned above, by checking the m^6^A-containing regions reported in lung adenocarcinoma cancer cell line A549 [81] and H1299 [82], we confirmed that no m^6^A peaks were further observed in A549 and H1299 for the variant-affected region (chr5:112176059-112176334, positive strand). Consequently, this LUAD somatic variant was upgraded to ‘medium’ confidence level in the m6A-CAVar database. Please note that, the predicted m^6^A dynamics in m6A-CAVar were systematically validated using available epitranscriptome datasets from the matched healthy and cancerous samples, providing another layer of quality assurance from real omics datasets: existing approaches only use those datasets to provide the m^6^A site information without searching for potential evidence of m^6^A status switching.

#### High confidence level

Only a very small number of variants have been clearly associated with diseases and phenotypes unveiled from GWAS analysis, and are known as disease-TagSNPs. These variants exhibited their clinical significance and are very likely to be functionally important. Thus, m^6^A variants of ‘high’ confidence level were defined as the validated m^6^A variants that can also be mapped to disease-TagSNPs extracted from ClinVar [71] and GWAS catalog [70]; while those not validated were referred to as “critical”.

Additionally, the association level (AL) between a SNP and m^6^A RNA modification was defined as following:

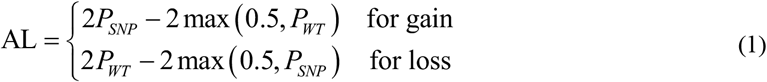

where, *P_WT_* and *P_SNP_* represent the probability of m^6^A RNA modification for the wild type and mutated sequences, respectively. The association level (AL) ranges from 0 to 1, with 1 indicating the maximum impact on m^6^A methylation. The statistical significance was assessed by comparing to the ALs of all mutations, with which the upper bound of the p-value can be calculated from its absolute ranking. The m^6^A-associated variants with association level > 0.4 and *P* < 0.1 were retained. We also considered the possibility of a variant destroying part of (but not an entire) m^6^A peak. For peaks wider than 500 nt, the impacts were also evaluated on the 200 nt flanking regions of the variant.

The predicted m^6^A variants were then validated on the epitranscriptome datasets from the matched health and cancer samples. We consider a prediction validated by omic data if the matched dynamics of m^6^A sites were observed under the heathy tissue and the cancer samples with the same tissue origin. It may be worth noting that, omic data was only used to inform the prediction of m^6^A sites in previous studies [55, 56, 79, 80]; however, our analysis also relies on it to confirm the predicted disturbance of m^6^A status between the health and cancer conditions. This extra layer of confirmation directly from available omic datasets should effectively enhance the reliability of our database.

### Functional annotation

The identified m^6^A variants were annotated with various information, including transcript region (CDS, 3’UTR, 5’UTR, start codon and stop codon), gene annotation (gene symbol, gene type, Ensembl gene ID), evolutionary conservation (phastCons 60-way), and deleterious level by SIFT [83], PolyPhen2 HVAR [84], PolyPhen2HDIV [84], LRT [85] and FATHMM [86] using the ANNOVAR package [87]. A total of 177,998 high-confidence m^6^A sites detected using base-resolution technology previously were collected and used to pin-point the precise location of the mediated m^6^A sites within the variant affected regions (**Sheet S3**). In addition, aspects of the post-transcriptional machinery that can be mediated by m^6^A methylation were also annotated including RBP binding regions from POSTAR2 [88], miRNA-RNA interaction from miRanda [89] and starBase2 [90], and splicing sites from UCSC [91] annotation with GT-AG role. Furthermore, to unveil potentially related pathogenesis, any association between disease and m^6^A variants was extracted from GWAS catalog [70] and ClinVar [71] databases.

### Database and web interface implementation

Hyper Text Markup Language (HTML), Cascading Style Sheets (CSS) and Hypertext Preprocessor (PHP) were applied to construct the m6A-CAVar web interface. All metadata was storage using MySQL tables. Besides, EChars was exploited to present statistical diagrams, and the Jbrowse genome browser [92] was included for interactive exploration and visualization of relevant records for genome regions of interests.

## Database content and usage

### Collection of m^6^A sites from 23 normal human tissues and 25 cancer cell lines in (m6A-TSDB)

In m6A-TSDB, a total of 184,554 and 499,369 m^6^A-containing peaks were collected from 23 normal human tissues and 25 cancer samples, respectively. Among them, 17 out of 25 tumor samples have the m^6^A profiles of their matched primary tissues. The m^6^A enriched peaks were called using exomePeak2 [76] with GC-correction function, after mapping the processed reads to human reference genome version hg19. It is worth mentioning that, for a more complete m^6^A epitranscriptome landscape view, a total of 177,998 base-resolution m^6^A sites collected from 27 datasets using six different m^6^A profiling techniques were integrated and used to pin-point the precise location of the mediated m^6^A sites within all tissue-specific m^6^A peaks (Sheet S3). All data collected in the m6A-TSDB can be freely downloaded or shared.

### Performance evaluation and model interpretation of tissue-specific m^6^A site prediction (m6A-TSFinder)

The performance of tissue-specific m^6^A site predictors was evaluated using 10-fold cross-validation and independent testing. For each distinct human tissue, we randomly selected 15% of experimentally validated m^6^A sites and used them as independent testing dataset. For 10-fold cross-validation, the training data was randomly divided into 10 sub-sections with the same number of positive and negative peaks. The prediction performance of each tissue-specific predictor was shown in Table 1. In general, the prediction accuracies for most tissues (20 out of the total 23 tissues) are in line with conventional approaches for m^6^A site prediction under strong supervision with base-resolution datasets, which typically reported a prediction performance between 0.8-0.85 in terms of AUROC [22, 93]. The performance for kidney (AUROC = 0.718), HSC (AUROC = 0.757) and brainstem (AUROC = 0.789) was somewhat worse, but the reasons are not very clear. Besides, in order to find the recurring sequence patterns preferred by each tissue-specific m6A prediction model, we further divided the peaks into instances of length (l=50) and extracted the consensus motifs from instances with predicted values higher than 0.5 using integrated gradient and TF-Modisco, under each tissue model, respectively. By trimming the overall letter frequencies with three gaps and two mismatches allowed, we identified one consistence motif under all tissue models (Figure S1) which was matched to the known m6A consensus motif DRACH. Please refer to Figure S1 for details.

**Table 1.** Performance evaluation of tissue-specific m^6^A model.

| Tissue type | 10-fold cross-validation |  |  |  | Independent testing |  |  |  |
| --- | --- | --- | --- | --- | --- | --- | --- | --- |
|  | Accuracy | Precision | MCC | AUROC | Accuracy | Precision | MCC | AUROC |
| Lung | 0.764 | 0.835 | 0.536 | 0.843 | 0.775 | 0.761 | 0.55 | 0.853 |
| Bladder | 0.758 | 0.760 | 0.517 | 0.836 | 0.766 | 0.750 | 0.532 | 0.848 |
| Colon | 0.740 | 0.770 | 0.482 | 0.810 | 0.744 | 0.730 | 0.490 | 0.810 |
| Lymph Nodes | 0.771 | 0.797 | 0.544 | 0.844 | 0.78 | 0.735 | 0.570 | 0.844 |
| Cerebrum | 0.745 | 0.799 | 0.495 | 0.827 | 0.758 | 0.768 | 0.515 | 0.834 |
| Cerebellum | 0.715 | 0.718 | 0.432 | 0.798 | 0.72 | 0.731 | 0.441 | 0.801 |
| Hypothalamus | 0.733 | 0.724 | 0.467 | 0.799 | 0.746 | 0.74 | 0.493 | 0.811 |
| Brainstem | 0.727 | 0.742 | 0.454 | 0.764 | 0.721 | 0.713 | 0.443 | 0.789 |
| Kidney | 0.685 | 0.694 | 0.369 | 0.755 | 0.647 | 0.628 | 0.297 | 0.718 |
| Bone Marrow | 0.694 | 0.634 | 0.391 | 0.757 | 0.698 | 0.721 | 0.397 | 0.757 |
| Liver | 0.742 | 0.747 | 0.484 | 0.805 | 0.737 | 0.717 | 0.476 | 0.803 |
| Ovary | 0.730 | 0.710 | 0.464 | 0.814 | 0.726 | 0.722 | 0.453 | 0.812 |
| Prostate | 0.752 | 0.779 | 0.507 | 0.819 | 0.759 | 0.736 | 0.521 | 0.830 |
| Soft Tissues | 0.766 | 0.855 | 0.544 | 0.855 | 0.771 | 0.775 | 0.543 | 0.858 |
| Skin | 0.750 | 0.850 | 0.511 | 0.835 | 0.773 | 0.753 | 0.547 | 0.857 |
| Stomach | 0.772 | 0.820 | 0.549 | 0.852 | 0.77 | 0.764 | 0.539 | 0.848 |
| Corpus Uteri | 0.722 | 0.656 | 0.452 | 0.813 | 0.734 | 0.715 | 0.470 | 0.822 |
| Adrenal Gland | 0.737 | 0.771 | 0.474 | 0.804 | 0.741 | 0.716 | 0.485 | 0.817 |
| Heart | 0.778 | 0.824 | 0.558 | 0.854 | 0.772 | 0.759 | 0.546 | 0.846 |
| Rectum | 0.747 | 0.725 | 0.496 | 0.826 | 0.767 | 0.747 | 0.536 | 0.828 |
| Testis | 0.743 | 0.770 | 0.489 | 0.810 | 0.731 | 0.734 | 0.463 | 0.804 |
| Thyroid Gland | 0.765 | 0.805 | 0.533 | 0.845 | 0.753 | 0.733 | 0.509 | 0.830 |
| Pancreas | 0.761 | 0.770 | 0.523 | 0.838 | 0.751 | 0.739 | 0.502 | 0.834 |

### Performance compared with existing approaches

We further compared the performance of the proposed m6A-TSFinder with existing m^6^A predictors specifically targeted at tissue level. *Dao et al*. previously developed an SVM-based model (iRNA-m6A) for m^6^A identification in human brain, liver, and kidney [65]. Later, im6A-TS-CNN [64] and TS-m6A-DL [66] further improved prediction performance by applying a convolutional neural network, using the same training and testing datasets provided in Dao’s work, respectively. It is worth mentioning that the training and testing datasets used in their work contain both 41nt-length positive and negative sequences with a m^6^A or non-m^6^A sites in the center. For a fair comparison, the same training and testing datasets were used to rebuild m6A-TSFinder in human brain, liver, and kidney, respectively. As described in the METHODS section, we applied a gated attention based multi-instance approach for identification of m^6^A signal in regions (~300nt). In this case, the 41nt-length sequences were treated as one instance and fed into the classifier, which makes the prediction performance comparable. As is shown in Table 2, when tested on independent dataset, m6A-TSFinder outperformed the three competing methods in two of the total 3 tissues tested (brain and liver), and achieved the best average performance (AUROC of 0.8593). The improvement may be due to the application of the LSTM layer after the convolutional layer, which enables the model to learn the long-range dependencies between the motifs. In addition, by learning from the low-resolution datasets, we expanded the human tissues supported from three to 23, which could significantly facilitate future research focusing on the dynamics of m6A methylome across different tissues.

**Table 2.** Performance comparison between m6A-TSFinder and competing approaches on independent dataset (AUROC)

|  | Performance on independent dataset |  |  |  |
| --- | --- | --- | --- | --- |
|  | m<sup>6</sup>A-TSFinder | TS-m<sup>6</sup>A-DL | im<sup>6</sup>A-TS-CNN | iRNA-m<sup>6</sup>A |
| Brain | 0.8132 | 0.8097 | 0.8056 | 0.7845 |
| Liver | 0.8850 | 0.8784 | 0.8805 | 0.8681 |
| Kidney | 0.8796 | 0.8802 | 0.8727 | 0.8565 |
| Average | 0.8593 | 0.8561 | 0.8529 | 0.8364 |

**Note:** For a fair comparison, the m<sup>6</sup>A-TSFinder was rebuilt for human brain, liver, and kidney, using the same training and testing datasets applied in the three previous works. The 41nt sequences were considered as one instance and fed into m<sup>6</sup>A-TSFinder.

### Assessing the impact of genetic variants on tissue-specific m^6^A sites by m6A-TSVar

The m6A-TSVar web server was designed to assess the impact of genetic variants on tissue-specific m^6^A RNA methylation using deep neural networks. The collected experimentally validated m^6^A peaks from 23 human tissues were integrated. The changes in the probability of m^6^A methylation affected by mutations were calculated, with the returned value of association level (AL) indicating how extreme the impact on m^6^A methylation is. To our best knowledge, the m6A-TSVar is the first web server for exploring m^6^A-affecting variants within a specific tissue by integrating the tissue-specific m^6^A patterns.

### Screening for cancer variants that affect m^6^A in their primary tissues (m6A-CAVar)

In m6A-CAVar, the cancer somatic variants from 27 TCGA projects were extracted. Their impacts on m^6^A RNA modification in the corresponding 23 healthy human tissues were evaluated and then systematically validated using 17 paired normal and tumor samples. A total of 587,983 cancer somatic variants were predicted to affect the m^6^A methylation status in their originating tissues (the ‘low’ confidence level group). Among them, the dynamic m^6^A status induced by 122,473 variants was observed on the available epitranscriptome profiles (the ‘medium’ confidence level group), and 1,718 confirmed m^6^A-variants were known to be associated with diseases and other phenotypes from GWAS analysis (the ‘high’ confidence level group) (see Table 3). Please refer to section of **Data collection and processing** for more details related to the classification of the m^6^A variants into different confidence group.

**Table 3.** Tissue-specific m^6^A cancer variants collected in m6A-CAVar.

| Cancer type | Primary tissue | Matched cancer cell line | Variant type | Classification |  |  | Total |
| --- | --- | --- | --- | --- | --- | --- | --- |
|  |  |  |  | Low | Middle | High |  |
| Lung Adenocarcinoma (TCGA-LUAD) | Lung | A549, H1299 | Gain | 27,845 | 6526 | 30 | 34,401 |
|  |  |  | Loss | 1233 | 1391 | 2 | 2626 |
| Bladder Urothelial Carcinoma (TCGA-BLCA) | Urinary Bladder | BCa5637 | Gain | 25,508 | 3702 | 13 | 29,223 |
|  |  |  | Loss | 3079 | 1691 | 6 | 4776 |
| Colon Adenocarcinoma (TCGA-COAD) | Colon | HT29, HCT116 | Gain | 30,540 | 8391 | 82 | 39,013 |
|  |  |  | Loss | 6 | 8284 | 74 | 8364 |
| Lymphoid Neoplasm Diffuse Large B-cell Lymphoma (TCGA-DLBC) | B Lymphocyte Cell Lines | OCI-Ly1 | Gain | 1189 | 82 | 2 | 1273 |
|  |  |  | Loss | 74 | 69 | 0 | 143 |
| Glioblastoma Multiforme (TCGA-GBM) | Cerebrum | U251, GOS-3, PBT003 | Gain | 8509 | 3648 | 47 | 12,204 |
|  |  |  | Loss | 1453 | 1181 | 12 | 2646 |
|  | Cerebellum |  | Gain | 8319 | 3659 | 38 | 12,016 |
|  |  |  | Loss | 1928 | 1271 | 4 | 3203 |
|  | Hypothalamus |  | Gain | 6723 | 3414 | 27 | 10,164 |
|  |  |  | Loss | 1522 | 1482 | 18 | 3022 |
|  | Brainstem |  | Gain | 7559 | 3168 | 40 | 10,767 |
|  |  |  | Loss | 1374 | 1451 | 8 | 2833 |
| Kidney Renal Clear Cell Carcinoma (TCGA-KIRC) | Kidney | iSLK.219 | Gain | 3844 | 227 | 4 | 4075 |
|  |  |  | Loss | 54 | 33 | 0 | 87 |
| Acute Myeloid Leukemia (TCGA-LAML) | Hematopoietic Stem Cells (HSC) | MOLM13, THP1, NOMO-1, MONO-MAC-6, MA9.3ITD | Gain | 448 | 274 | 0 | 722 |
|  |  |  | Loss | 3 | 35 | 3 | 41 |
| Liver Hepatocellular Carcinoma (TCGA-LIHC) | Liver | HepG2, Huh7, SMMC7721, HCCLM3 | Gain | 7416 | 2511 | 2 | 9929 |
|  |  |  | Loss | 18 | 1765 | 4 | 1787 |
| Ovarian Serous Cystadenocarcinoma (TCGA-OV) | Ovary | PEO1 | Gain | 7022 | 531 | 0 | 7553 |
|  |  |  | Loss | 1350 | 1090 | 6 | 2446 |
| Prostate Adenocarcinoma (TCGA-PRAD) | Prostate Gland | Cd-RWPE-1 | Gain | 3825 | 636 | 6 | 4467 |
|  |  |  | Loss | 550 | 288 | 2 | 840 |
| Sarcoma (TCGA-SARC) | Soft Tissues | U20S | Gain | 3592 | 1324 | 4 | 4920 |
|  |  |  | Loss | 373 | 28 | 0 | 401 |
| Skin Cutaneous Melanoma (TCGA-SKCM) | Skin | Mel624 | Gain | 79,470 | 17,177 | 118 | 96,765 |
|  |  |  | Loss | 6472 | 1559 | 2 | 8033 |
| Stomach Adenocarcinoma (TCGA-STAD) | Stomach | BGC823 | Gain | 35,438 | 2202 | 34 | 37,674 |
|  |  |  | Loss | 1103 | 3313 | 27 | 4443 |
| Uterine Corpus Endometrial Carcinoma (TCGA-UCEC) | Corpus Uteri | HEC-1-A | Gain | 80,712 | 38,242 | 266 | 119,220 |
|  |  |  | Loss | 7813 | 1828 | 22 | 9663 |
| Lung Squamous Cell Carcinoma (TCGA-LUSC) | Lung | - | Gain | 31,106 | - | 118 | 31,224 |
|  |  |  | Loss | 2328 | - | 2 | 2330 |
| Mesothelioma (TCGA-MESO) | Lung | - | Gain | 595 | - | 4 | 599 |
|  |  |  | Loss | 57 | - | 0 | 57 |
|  | Heart | - | Gain | 674 | - | 5 | 679 |
|  |  |  | Loss | 102 | - | 0 | 102 |
| Brain Lower Grade Glioma (TCGA-LGG) | Cerebrum | - | Gain | 6714 | - | 92 | 6806 |
|  |  |  | Loss | 1423 | - | 19 | 1442 |
|  | Cerebellum | - | Gain | 6601 | - | 109 | 6710 |
|  |  |  | Loss | 1745 | - | 16 | 1761 |
|  | Hypothalamus | - | Gain | 5010 | - | 77 | 5087 |
|  |  |  | Loss | 1698 | - | 11 | 1709 |
|  | Brainstem | - | Gain | 5740 | - | 114 | 5854 |
|  |  |  | Loss | 1528 | - | 13 | 1541 |
| Kidney Chromophobe (TCGA-KICH) | Kidney | - | Gain | 484 | - | 9 | 493 |
|  |  |  | Loss | 9 | - | 0 | 9 |
| Kidney Renal Papillary Cell Carcinoma (TCGA-KIRP) | Kidney | - | Gain | 4028 | - | 17 | 4045 |
|  |  |  | Loss | 118 | - | 0 | 118 |
| Cholangiocarcinoma (TCGA-CHOL) | Liver | - | Gain | 728 | - | 2 | 730 |
|  |  |  | Loss | 166 | - | 2 | 168 |
| Adrenocortical Carcinoma (TCGA-ACC) | Adrenal Gland | - | Gain | 2285 | - | 21 | 2306 |
|  |  |  | Loss | 385 | - | 3 | 388 |
| Pheochromocytoma and Paraganglioma (TCGA-PCPG) | Adrenal Gland | - | Gain | 345 | - | 1 | 346 |
|  |  |  | Loss | 57 | - | 0 | 57 |
| Rectum Adenocarcinoma (TCGA-READ) | Rectum | - | Gain | 12,433 | - | 100 | 12,533 |
|  |  |  | Loss | 1098 | - | 4 | 1102 |
| Thymoma (TCGA-THYM) | Heart | - | Gain | 520 | - | 7 | 527 |
|  |  |  | Loss | 80 | - | 2 | 82 |
| Testicular Germ Cell Tumors (TCGA-TGCT) | Testis | - | Gain | 405 | - | 3 | 408 |
|  |  |  | Loss | 109 | - | 0 | 109 |
| Thyroid Carcinoma (TCGA-THCA) | Thyroid Gland | - | Gain | 992 | - | 6 | 998 |
|  |  |  | Loss | 156 | - | 0 | 156 |
| Pancreatic Adenocarcinoma (TCGA-PAAD) | Pancreas | - | Gain | 6473 | - | 55 | 6528 |
|  |  |  | Loss | 1236 | - | 3 | 1239 |
| **Total** | - | - | - | 463,792 | 122,473 | 1718 | 587,983 |

### Deciphering the tissue-specificity of cancer m^6^A variants

Of interest is whether m^6^A variants function in different cancer growing tissues. For this purpose, we calculated the proportion of m^6^A variants that function in different numbers of tissues, and the results suggested that most m^6^A-associated cancer variants are tissue- and cancer-specific (93.25%), while only around 1.17% are functional in the originating tissues of more than three types of cancers (Figure 4A). The consistency is much higher at gene level. Only around 16.59% of m^6^A variant carrying genes are associated with a single tissue. More than 60.29% were shared in more than three tissue types (Figure 4B), suggesting some common epitranscriptome layer circuitry at the gene level in different cancers. We further examined the proportion of shared m^6^A variant-carrying genes between two different tissues. As shown in Figure 4C, most tissues eg. skin and stomach have a strong correlation with each other. However, tissues like heart, testis and thyroid showed rather weak association with other tissues, which may suggest more tissue-specific epitranscriptome circuitry for cancers originating in those tissues.

**Figure 4.**
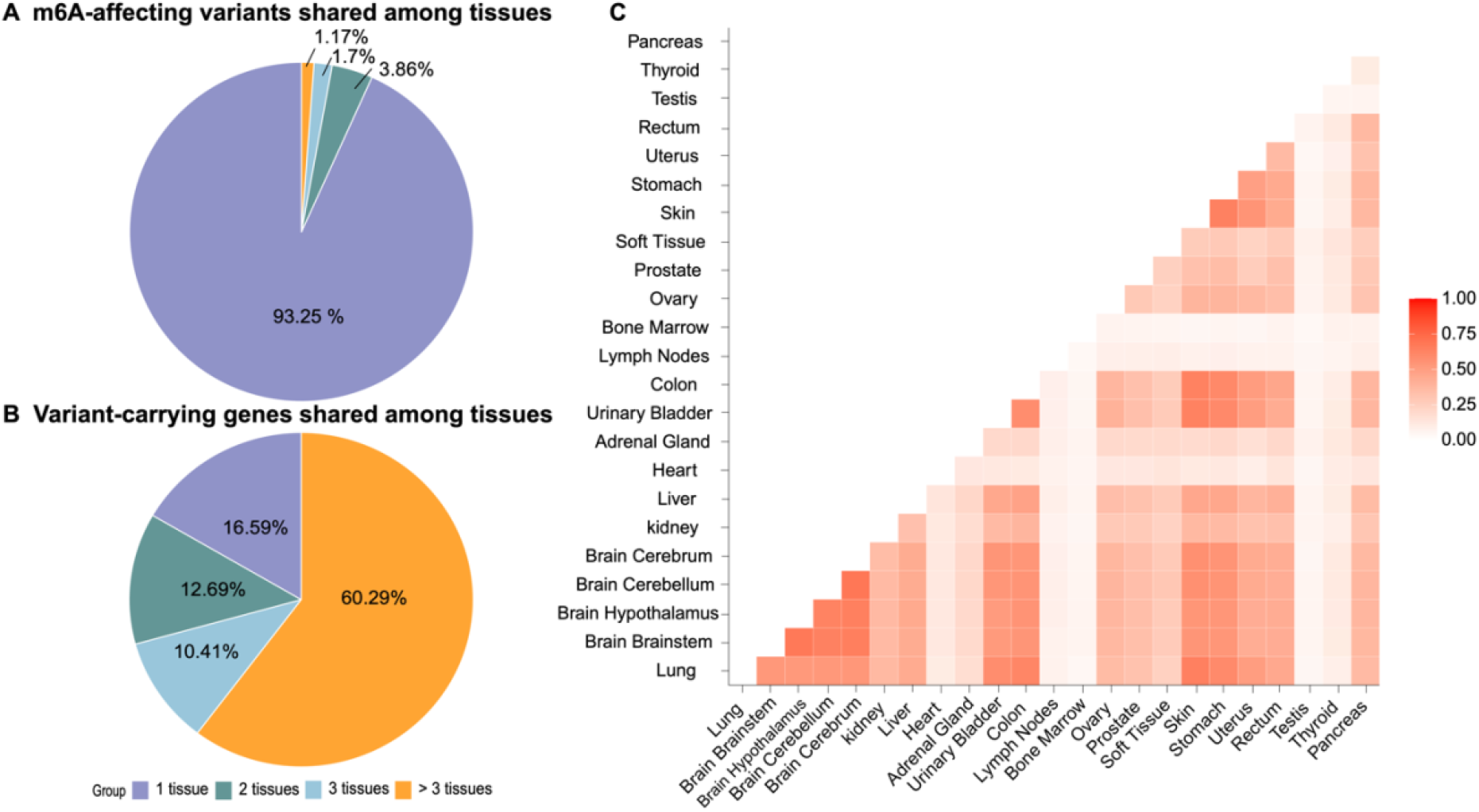
Tissue-specificity of cancer m^6^A variants. **(A)** The proportion of m^6^A variants that are shared among different tissues. Most m^6^A-associated variants (93.25%) were identified in only 1 tissue, with 3.86%, 1.7%, and 1.17% identified in 2, 3 and more than 3 tissues, respectively. (**B**) The proportion of m^6^A variant-carrying genes shared among tissues, the consistency is much higher at gene level. Most m^6^A variants-carrying genes are shared among multiple tissues, with only 16.59% being associated to one tissue type. **(C)** The pairwise association of tissues in terms of proportion of shared m^6^A variant carrying genes. Most tissues are significantly correlated. The exceptions are heart, adrenal gland, lymph nodes, bone marrow, testis and thyroid.

We finally identified the m^6^A variant-carrying genes that are associated with the most TCGA cancer types. Only experimentally validated m^6^A variants (medium confidence level and above) were considered here for a more reliable analysis. Top of the list was CENPF where variants may change its m^6^A methylation status in the primary tissue of 15 cancer types, followed by DST, MKI67 and PLEC, which were all related to 14 cancer types (detailed in Sheet S4). Among them, the roles in epitranscriptome regulation of CENPF, MKI67 and PLEC have been indicated previously in glioblastoma [94], breast cancer [95] and pancreatic cancer [96], respectively.

### Enhanced web interface and application

The m6A-TSHub features a user-friendly web interface with multiple useful functions, including databases and online-servers, which enable users to fast query databases, upload their own custom jobs, and download all m^6^A-related information at tissue level. The collected functional m^6^A-affecting variants can be queried using a human body diagram according to their primary tissues (Figure 5A), as well as by different cancer types along with further filters (e.g., gene type, m^6^A status, confidence level, and disease association, Figure 5B). The query function also returns several categories of useful information, including TCGA project names [77], tumor-growth tissues, genes, chromosome regions, COSMIC ID [97], and disease phenotypes (Figure 5C). The details of tissue-specific m^6^A peaks collected in m6A-TSDB (Figure 5D) and cancer m^6^A-associated variants in m6A-CAVar (Figure 5E) can be viewed by clicking the site or variant ID, along with annotated disease-association regulations (Figure 5F). Furthermore, online servers allow for identification of m^6^A sites and m^6^A-associated variants within user-defined regions, with 23 types of human tissues to be selected (Figure 5G-H). A genome browser is available for interactive exploration of the genome regions of interest. All metadata provided in the m6A-TSHub can be freely downloaded. Users can refer to the ‘Help’ page for more detailed guidance and instructions.

**Figure 5.**
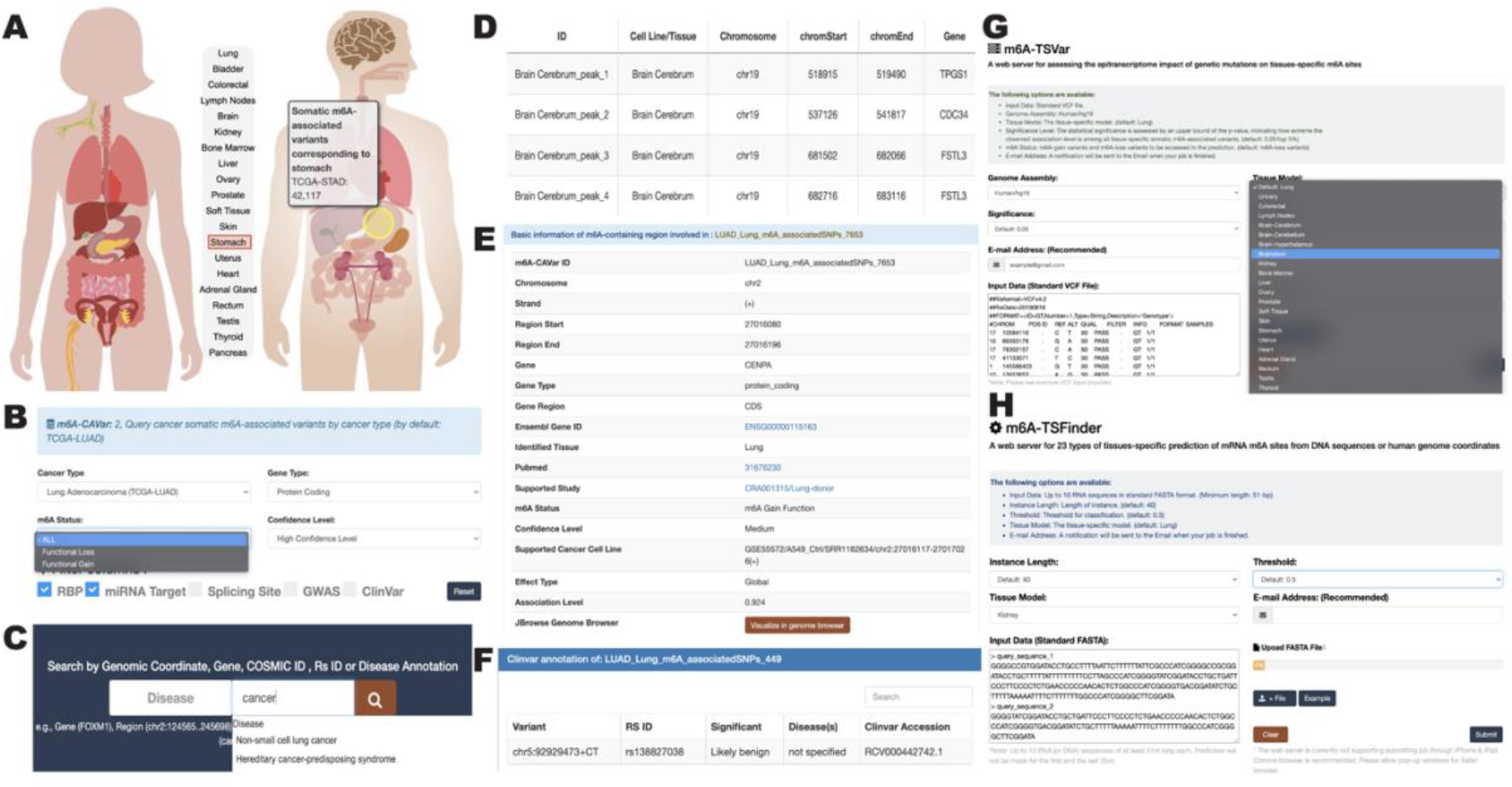
Enhanced web interface. **(A)** A human body diagram is available for querying cancer somatic m^6^A-associated variants in their originating tissues. **(B-C)** Users can also query the associated variants by cancer type, disease, region, gene symbol, COSMIC and rs ID, and further filter the returned results. (**D**) Details of tissue-specific m^6^A peaks collected in m6A-TSDB. **(E-F)** Details of cancer-related m^6^A-associated variants and disease annotation involved. **(G-H)** The online-tools provided for analysis of user-uploaded files, including assessing m^6^A-associated variants in tissues (m6A-TSVar) and identifying tissue-specific m^6^A sites (m6A-TSFinder), respectively.

### Utility case study 1: PIK3CA variant in colon cancer

Previous studies have reported that m^6^A RNA modification plays an important role in colon cancer [61, 98–100]. The TCGA-COAD project [77] presented a large number of somatic variants identified from various colon adenocarcinoma samples. However, it is still unclear which single genetic variant may lead to m^6^A dysregulation. In m6A-CAVar, a somatic variant at chr3: 178952085 (A>T) on PIK3CA identified from TCGA-COAD project (TCGA barcode: TCGA-AA3821-01A-01W-0995-10) was predicted to erase the m^6^A methylation of a region (chr3: 178951888-178952363, positive strand). The m^6^A methylation was observed in human healthy colon, but disappeared in the colon adenocarcinoma cancer cell line HCT116 [101]. This somatic variant is also recorded in the COSMIC database from colon tumor samples under the legacy identifier of COSM776, and reported to be associated with 27 submitted interpretations and evidences in the ClinVar database [71], including PIK3CA related overgrowth spectrum (ClinVar accession: RCV000201235.1), breast adenocarcinoma (ClinVar accession: RCV000014629.5), and pancreatic adenocarcinoma (ClinVar accession: RCV000417557.1). Taken together these observations strongly support the functional importance of this variant. Additionally, the m^6^A-associated variant falls within the binding regions of two RNA binding proteins (TARDBP and NUDT21), whose interaction may be regulated by the loss of m^6^A methylation in the cancer condition, providing some putative downstream regulatory consequences of the variant.

### Utility case study 2: PLEC variant in glioblastoma

Glioblastoma (GBM) is the most aggressive type of brain tumor and is associated with rising mortality. The roles of m^6^A regulators in this disease have been previous indicated [102–105]. A somatic cancer variant on PLEC was identified from the TCGA-GBM project (TCGA barcode: TCGA-06-5416-01A-01D-1486-08) at chr8: 144991388 (C>T). This cancer variant was predicted to lead to gain of a m^6^A site on a previously un-methylated region in healthy human cerebrum. Indeed, an m6A site was detected at this region from malignant glioblastoma tumor cell line U-251. This mutation has a record in ClinVar database (ClinVar accession: RCV000177727.1). Screening for potential post-transcriptional regulations revealed that the cancer variant falls within the target binding regions of six RNA binding proteins, including the m^6^A reader YTHDF1, which are known to bind m^6^A-containingRNAs and promote cancer stem cell properties of glioblastoma cells [106]. It should be of immediate interest to ask whether the methylation of PLEC regulates its interaction with YTHDF1 and other RBPs, and what the functional consequences are.

### Utility case study 3: EGFR variant in lung cancer

The associations between m^6^A RNA modification and human lung cancers have been well studied. The m^6^A eraser FTO may be a prognostic factor in lung squamous cell carcinoma (TCGA-LUSC) [107], and the m^6^A writer METTL3 regulates EGFR expression to promote cell invasion of human lung cancer cells [82]. The m6A-CAVar database can be used to explore the role of m^6^A variants of EGFR in lung cancers. We first search by gene name ‘EGFR’ at the front page of m6A-CAVar database, then filter the results and keep only records related to lung tissue, which retains a total of 10 cancer m^6^A-associated variants from two lung cancer types (Figure 6A-B). Alternatively, the users can query all recorded m^6^A-associated variants that functions in lung tissue by simply clicking the relevant part from human body diagram (Figure 6C). More details can be accessed by clicking the variant ID. For example, if we check further the details of a m^6^A-gain variant from TCGA-LUAD project at chr7: 55259515 (T>G), we can see that this variant was recorded in the ClinVar database and is relevant to eight disease conditions including lung cancers (Figure 6D), which may suggest potential cancer pathogenesis originating in the epitranscriptome layer.

**Figure 6.**
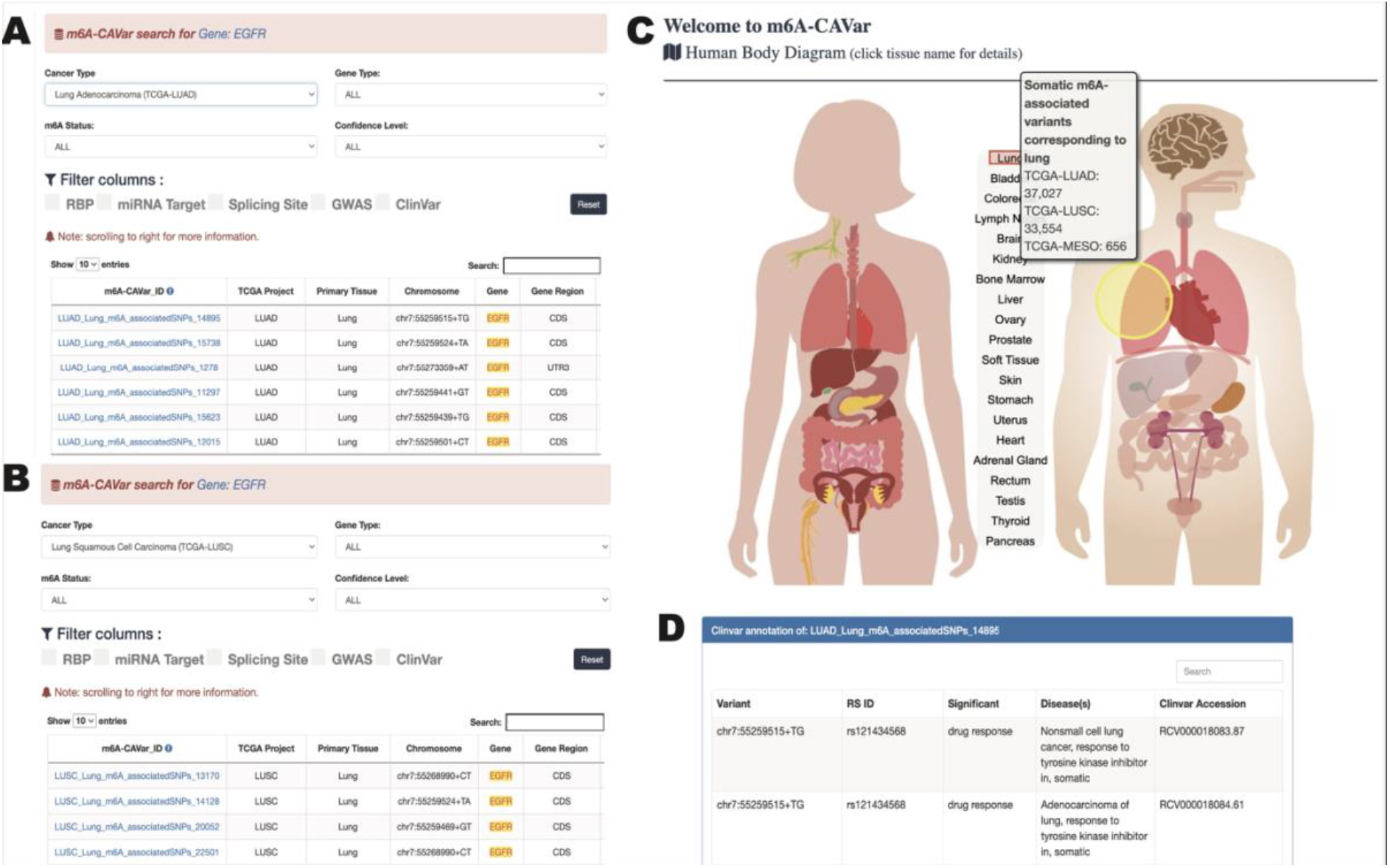
Case study on gene EGFR. **(A and B)** Searching for the gene ‘EGFR’ in m6A-CAVar database returns a total of 10 m^6^A variants, the details of which can be viewed by clicking the m6A-CAVar ID. **(C)** A human body map is provided at front page of m6A-CAVar website, which enables quick positioning of cancer m^6^A-associated variants functions at a specific tissue. **(D)** The disease and phenotype association of recorded m^6^A variant.

## Discussion and perspectives

The context-specific expressions and functions of m^6^A regulations have been repeatedly reported in existing studies [59–63], suggesting the involvement of the tissue-specific m^6^A methylome in essential biological processes and multiple disease mechanisms. Besides, the associations between RNA methylation levels and the activities of RNA methylation regulators were clearly unveiled, reporting that there exist some condition-specific RNA co-methylation patterns (a group of RNA m6A methylation sites whose methylation levels go up and down together) [108–110]. These co-methylation patterns are enriched by the substrate targets of m6A regulators, and thus are probably regulated by specific m6A methyltransferase or demethylase.

Here we present m6A-TSHub, a comprehensive online platform for unveiling the context-specific m^6^A methylation and m^6^A-affecting mutations in 23 human tissues and 25 tumor conditions. In m^6^ATSHub, a total of 184,554 and 499,369 m^6^A sites derived from 23 human normal tissues and 25 matched tumor samples were collected (m6A-TSDB), from which some potential patterns for the tissue specific m6A modification sites were revealed (e.g., heart-enriched gene RYR2 and PXDNL, see Figure S2). Based on these collected data, 23 distinct m^6^A prediction models were built in tissue level, using deep neural networks (m6A-TSFinder). In addition, to elucidate the genetic factor of epitranscriptome dysregulation, m6A-CAVar identified a total of 587,983 cancer somatic mutations that may alter the m^6^A status in the corresponding cancer originating tissues, and annotated them with various functional annotations, including features relating to post-transcriptional regulations (RBP binding regions, microRNA targets, splicing sites), disease and phenotype association, as well as other useful genomic information (transcript structure, phastCons, deleterious level) to provide a more comprehensive overview. We also provided a web server m6A-TSVar for assessing the effect of genetic variants on m6A methylation in a specific tissue.

While most of existing approaches for RNA modification site prediction ignore the tissue-specific signatures of m^6^A methylation, by taking advantage of existing tissue-specific epitranscriptome data our method can predict the m^6^A methylation within a specific tissue. Compared with exiting approaches for tissue-specific m^6^A methylation site prediction [64–66], our approach m6A-TSFinder achieved a higher prediction performance (see **Table 2**) and hugely expanded the number of supported tissue types from 3 to 23 (see Table 1).

Compared with existing approaches for decoding the epitranscriptome impact of genetic variants, m6A-CAVar has the following two major advantages. First, m6A-CAVar relies on a finer prediction model (m6A-TSFinder) that appreciates the specific pattern of RNA methylomes across different tissues. By directly learning from the epitranscriptome profiles in 23 healthy human tissues, m6A-CAVar was able to evaluate the tissue-specific impact of cancer somatic variants on m^6^A modification in their originating tissue, providing a more detailed picture of the genome-epitranscriptome association. This improves on existing approaches that ignore the distinct signatures of RNA methylation across different tissues and thus failed to address tissue-specific effects. Second, the predicted m^6^A dynamics in m6A-CAVar were systematically validated using available epitranscriptome datasets from the matched healthy and cancerous samples, providing another layer of quality assurance from real omics datasets. In contrast, existing approaches use those datasets only to provide the m^6^A site information without searching for potential evidence of m^6^A status switching.

To date, epitranscriptome data is still rather scarce. Due to the limited availability of datasets, matched healthy tissue and cancer m^6^A profiling samples are only available for 14 out of the total 27 cancer types, prohibiting a more thorough validation of the predicted results. Furthermore, substantial discrepancy has been observed among different RNA modification profiling approaches that can capture different technical bias [111–114], which can produce additional inaccuracy. Currently, context-specific epitranscriptome prediction is only possible for a small number of conditions (cell line, tissue type, treatment) with data [64–66]. However, the m6A-TSHub framework will be further expanded when epitranscriptome datasets are more abundantly available for a more comprehensive and less biased screening of context-specific m^6^A-variants, along with linking the tissue-specific epitranscriptome patterns with other important cancer-associated factors such as human aging [67, 115]. Particularly promising is the recent development in Nanopore direct RNA sequencing technology that enables simultaneous identification of multiple RNA modifications with simplified sample preparation procedures [116–124].

## Data availability

The data underlying this article are available via www.xjtlu.edu.cn/biologicalsciences/m6ats, and in its online supplementary material. The online version of m6A-TSFinder and m6A-TSVar web server are available via www.xjtlu.edu.cn/biologicalsciences/m6ats by clicking ‘Tool’ section. The local version and project codes can be accessed on the ‘Download’ page.

## CRediT author statement

**Bowen Song:** Methodology, Data Curation, Software, Visualization, Writing-Original Draft. **Daiyun Huang**: Software, Supervision. **Yuxin Zhang**: Visualization. **Zhen Wei**: Resources, **Jionglong Su**: Visualization. **João Pedro de Magalhães**: Visualization. **Daniel J. Rigden**: Writing-Review & Editing. **Jia Meng**: Conceptualization, Supervision, Writing-Review and Editing, Funding acquisition. **Kunqi Chen**: Conceptualization, Resources, Writing-Review and Editing, Funding acquisition.

## Acknowledgements

This work was supported by the National Natural Science Foundation of China (Grant No. 32100519 and 31671373) and XJTLU Key Program Special Fund (Grant No. KSF-T-01, KSF-E-51, and KSF-P-02). The authors also acknowledge the researcher of the various resources mentioned in the manuscript to share their data, especially for tissue-specific m^6^A sequencing data.

